# Traumatic brain injury has a lasting impact on hippocampal neurogenesis and Notch1 is involved in regulating this injury response

**DOI:** 10.64898/2026.02.03.703567

**Authors:** Nicole M. Weston, Timothy N. Keoprasert, Jakob C. Green, Sarah Baig, Dong Sun

## Abstract

Traumatic brain injury (TBI) induces a series of neuropathological changes in the brain including neurogenesis, an important cellular response involved in brain repair and regeneration. TBI-enhanced neurogenesis in the dentate gyrus (DG) of the hippocampus is of particular importance due its contribution to learning and memory functions. In the neurogenic process, proliferation and differentiation of neural stem cells (NSCs) follow a well-characterized sequence controlled by many factors including Notch1, which plays essential roles in regulating NSC fate determination under physiological conditions in both developing and adult brains. Following TBI, the dynamic changes of NSCs and the involvement of Notch1 on their development at different stages post-injury are not fully characterized. In the current study, we examined the impact of TBI and Notch1 on NSCs proliferation, survival and neuronal differentiation. Utilizing transgenic mice with tamoxifen-induced GFP expression and Notch1 knock-out in nestin+ NSCs, we examined DG neurogenic response at acute, subacute and chronic stages following a moderate lateral fluid percussion injury. We found that TBI enhanced a proliferative response in the DG at the acute stage following injury; however, this injury response was abolished when Notch1 was conditionally deleted from nestin+ NSCs. We also found that injury and Notch1 deletion drove NSCs committing fate choice towards neuronal differentiation. The results of this study provides further knowledge regarding TBI-induced neurogenic response and Notch1 as the key regulating mechanism.

## 1. Introduction

Traumatic brain injury (TBI) induces a host of pathophysiologic responses including enhanced endogenous neurogenic response in the primary neurogenic regions, i.e., the subventricular zone (SVZ) and dentate gyrus (DG) of the hippocampus (Eriksson et al. 1998; Gage and Temple 2013). Under normal homeostatic conditions, in the adult brain, these neurogenic regions continuously supply new neurons that mature into functional neurons with cells generated from the DG becoming DG granular neurons involved in hippocampal-dependent learning and memory functions (Eriksson et al. 1998; Anacker and Hen 2017; Aimone, Deng, and Gage 2011; Deng, Aimone, and Gage 2010; Marin-Burgin and Schinder 2012). Following TBI, the fate and functional contribution of injury-induced new neurons in the hippocampus are not fully understood. It is postulated that this injury-induced response is likely associated to the post-TBI endogenous repair process (Chirumamilla et al. 2002; Blaiss et al. 2011; Sun et al. 2007; Sun et al. 2015).

Thus far, the TBI-induced neurogenic response has been explored in various models of TBI. Across all models, studies have consistently shown an injury-induced proliferative response in neurogenic regions of the adult brain (Chirumamilla et al. 2002; Villasana et al. 2015; Sun 2016; Bye et al. 2011; Redell et al. 2020). This cellular response has a strong time dependency occurring in the first week after TBI with 2 days post-injury as the peak time (Dash, Mach, and Moore 2001; Rice et al. 2003; Sun et al. 2005). As the hippocampus is vulnerable to TBI, and hippocampal neurogenesis plays an important role in learning and memory functions, the injury-induced neurogenic response in the hippocampus is heavily studied. Thus far, the fate and functions of the injury-induced proliferative populations in the hippocampus have been explored. Studies have demonstrated that many of the injury-enhanced proliferative cells go through apoptosis within the first 4 weeks after generation, while those that survived matured into functional neurons participating hippocampal dependant learning and memory functions (Sun et al. 2007; Blaiss et al. 2011; Sun et al. 2015; Weston et al. 2021; Weston et al. 2024). Furthermore, injury-induced DG new neurons has shown biphasic responses in excitatory activity at different stages following TBI demonstrating their involvement in dynamic neuronal network remodeling in the DG (Danielewicz et al. 2025). These findings established the importance of this injury-induced cellular response for the endogenous repair and regeneration of the injured brain.

The involvement of the neurogenic response in post-TBI functional recovery suggests the potential of developing stem cell therapies for brain regeneration by harnessing this response (Weston and Sun 2018). Towards this goal, a better understanding of the mechanisms that regulate the injury-induced neural stem cell (NSC) response is necessary. Among possible mechanisms, Notch1 is a strong contender due to its widely essential roles in NSC maintenance, proliferation, and survival in both development and adult neurogenic processes (Androutsellis-Theotokis et al. 2006; Imayoshi et al. 2010; Ding et al. 2016; Ables et al. 2010). Under physiological conditions, Notch1 signaling controls the number of cells remaining in the NSC pool or exiting this reserve through communication with neighboring cells (Ables et al. 2010; Imayoshi et al. 2010; Aguirre, Rubio, and Gallo 2010). It is unknown if Notch1 is also an essential driver controlling the injury-induced neurogenic response. In a rat focal cerebral ischemic injury model, Notch1 and its signaling pathway proteins have shown increased expression in the SVZ strongly associated to injury-induced cell proliferation (Wang et al. 2009; Sun et al. 2013). Thus, it is likely that Notch1 signaling is involved in the NSC response following brain injury. In this study, we will ascertain the role of Notch1 in regulating the TBI-induced neurogenic response in the hippocampus.

As Notch1 is involved in cell proliferation in many types of cells, we generated a transgenic mouse line with conditional Notch1 knockout in nestin+ cells allowing us to directly investigate the importance of Notch1 in regulating the post-injury NSC response. Utilizing transgenic mouse lines with tamoxifen (TAM) induced eYFP expression and Notch1 gene cKO in nestin+ NSCs, in addition with thymidine analog bromodeoxyuridine (BrdU) to tag proliferating NSCs at different time point post-TBI, we examined the impact of TBI on NSC proliferation, survival and neuronal differentiation at the acute, subacute and chronic stages following TBI. We also investigated the consequences of Notch1 conditional knock out in nestin+ NSCs on this injury-induced DG neurogenic response.

## 2. Materials and Methods

### 2.1. Experimental Animals

A mixed group of female and male adult mice at 8 to 10 weeks old were used. A total number of 72 animals were included (n = 4-8/group) in this study. These mice were derived from cross-breeding of three strains (Jackson Laboratories) by the VCU Transgenic Core to create a control line (NC-Y) and a Notch1 conditional knockout line (NC-NKO-Y, Notch 1cKO) (**Figure 1a**). The nestin-CreER (Jax. Stock No: 012906) and R26R-EYFP (Jax. Stock No: 006148) strains were used to produce the NC-Y control animals, and the Notch1flox (Jax. Stock. No: 007181) strain was added to produce the NC-NKO-Y Notch1 cKO animals. These lines of mice have been well characterized previously (Ables et al. 2010). Tails snips were sent to Transnetyx for genotyping to confirm the Notch1 cKO and strains were maintained as homozygotes. The mice were housed in a facility with a 12hr light-dark cycle, controlled humidity and temperature, and food and water were provided ad-libitum. All procedures were approved by the VCU Institutional Animal Care and Use Committee under the protocol of AM10222. In studies examining neurogenesis, there were scattered reports about sex or estrogen transiently affecting baseline cell proliferation in the DG in rats (Tanapat et al. 1999; Yagi et al. 2020), no significant sex differences were found affecting long term adult neurogenesis in rodents (Yagi and Galea 2019). In this study, no sex-related difference was found thus data from males and females were combined.

**Figure 1.**
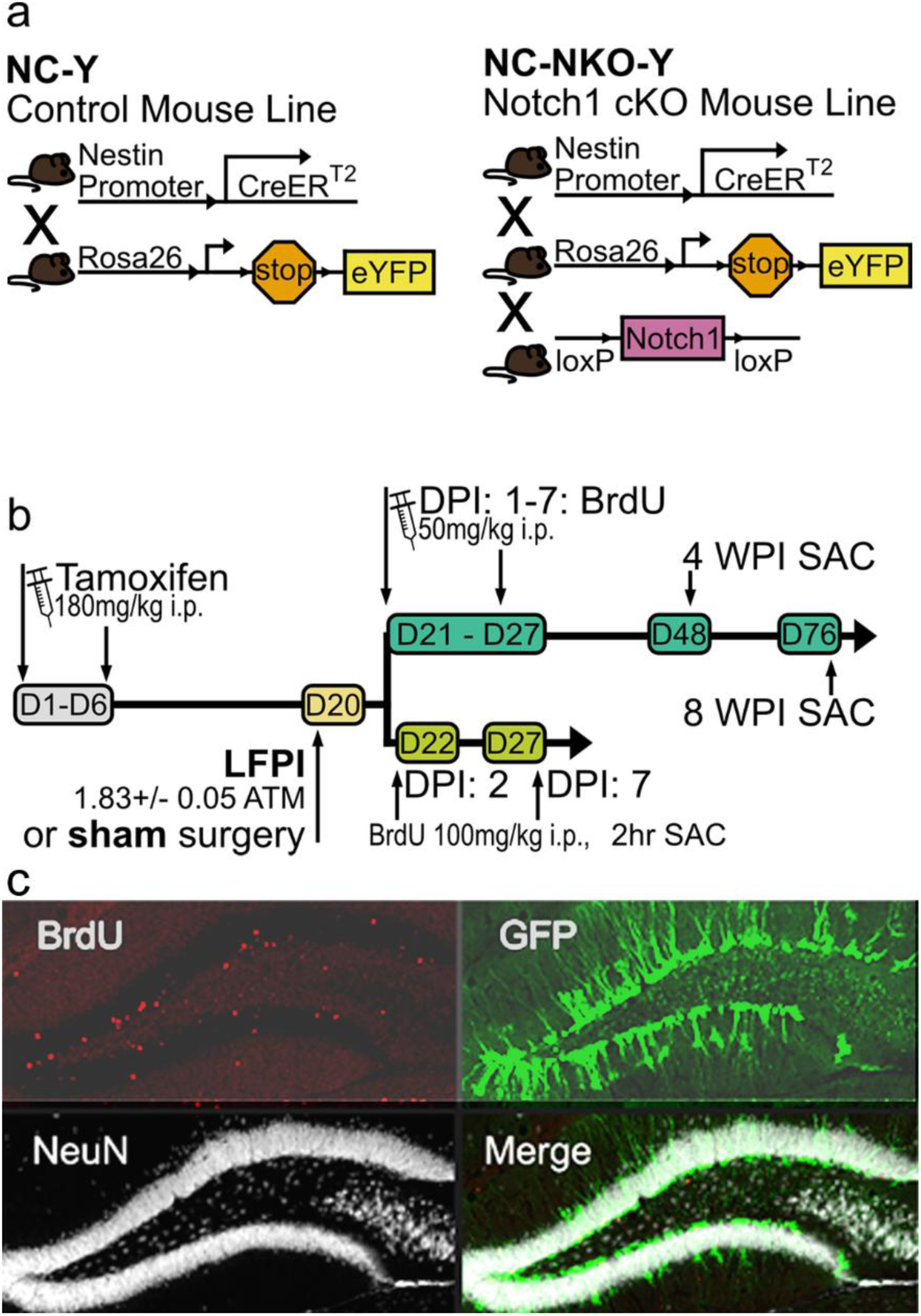
Transgenic mouse lines and experiment timeline. (**a**) The two transgenic mouse lines used for experiments. NC-Y is the control line comprised of the separate strains nestin-CreER and R26R-EYFP. NC-NKO-Y is the Notch1 cKO line comprised of nestin-CreER, R26R-EYFP, and Notch1flox. (**b**) Experimental timeline. Animals received 6 i.p. injections of tamoxifen followed by injury or sham surgery 2 weeks post-tamoxifen. Animals were then divided into two separate groups. On either day 2 or day 7 post-injury, the first group received a single BrdU i.p injection (100mg/kg) and were sacrificed two hours later. The second group received BrdU i.p. injections (50mg/kg) every day for the first 7 days post-injury and were sacrificed 4 or 8 weeks post-injury. (c) Representative confocal images of BrdU+, GFP+ and NeuN+ cells taken on a LSM710 confocal with a 20x objective.

To induce Cre expression in the transgenic mouse models, tamoxifen (TAM) was administrated i.p. (180mg/kg in 10% EtOH/sunflower oil) single daily for six consecutive days (**Figure 1b**).

### 2.2. Surgical Procedure

Two weeks after the last TAM administration animals were randomly divided into groups receiving either a moderate lateral fluid percussion injury (LFPI) or sham surgery. Briefly, the animal was placed in an acrylic chamber with 4% isoflurane for anesthesia induction and then fixed on a stereotaxic frame under a continuous flow of 2.5% isoflurane in O_2_ through a nosepiece. After a midline incision to expose the skull, a craniotomy was made using a 2.7mm trephine at the left parieto-temporal bone halfway between lambda and bregma. A Luer-slip hub made with a 20-gauge needle cap was placed at the craniotomy site with cyanoacrylate and dental acrylic. Anesthesia was turned off and the animal was placed in a heated cage to recover from surgical procedure. Two hours later, the animal was re-anesthetized with 4% isoflurane. The hub placed over the craniotomy site was filled with 0.9% saline and attached to a recalibrated FPI device to administer a pulse target of 1.83±0.05ATM inducing a moderate TBI. Immediately following injury, the animal was placed on a heated pad and righting time was recorded to further confirm the injury severity. Once righted, the hub was removed and the skin incision sutured, the animal was returned to the heated cage for recovery before being returned to the vivarium. Animals in the sham group went through the same surgical procedure without receiving the fluid pulse. All animals received post-operative care and hydrogel supplement was added to their home cages to aid in hydration.

### 2.3. BrdU administration

After injury, animals were randomly divided into 4 study groups according to sacrificing time point: 2- or 7-days post-injury (DPI), or 4- or 8-weeks post-injury (WPI) (**Figure 1b**). The 2- and 7-DPI animals received a single pulse BrdU administration i.p. (100mg/kg) at 2 hours before perfusion to study injury-induced cell proliferation at 2- and 7-DPI, whereas mice in the 4- and 8-WPI groups received i.p. BrdU single daily injection (50mg/kg) at 1-7 DPI to study the survival of injury-induced newly generated cells.

### 2.4. Immunohistochemistry

Mice were sacrificed with transcardiac perfusion with 4% paraformaldehyde followed by PBS at the designated post-injury time point described above. Brains were dissected, post-fixed, 50µm thick coronal sections were sliced with a vibratome and collected in a 48-well plate containing PBS plus 0.01% sodium azide. To examine injury-induced cell proliferation (2- and 7-DPI groups), consecutive five sections 400µm apart spanning a 2000µm thickness of the DG were processed for BrdU chromogenic labeling. To study cell survival (4- and 8-WPI groups), within a span of 1000µm of the DG, every 5^th^ section was processed for a total of 4 sections per animal.

Sections were washed with PBS followed by application of a blocking solution and sequentially the primary antibody solution. For the 2- and 7-DPI groups, the primary antibody used was rat monoclonal anti-BrdU (1:2000; Abcam AB6236) and secondary antibody was biotinylated anti-rat IgG (1:200; Vector Laboratories BA-9401), and the liquid form of 3,3’-Diaminobenzidine (DAB) was used as the chromogenic. For the 4- and 8-WPI groups, the primary antibodies used were rabbit anti-GFP (1:2000; Invitrogen A11122), mouse anti-NeuN (1:100; Millipore MAB377) and the rat anti-BrdU. The primary antibody anti-GFP was used to target the transgenically expressed eYFP, as YFP and GFP have a very similar peptide sequence homology. As a result, we will refer to staining patterns using GFP+ and eYFP when referring to the mouse strain. Secondary antibodies used were biotinylated anti-rabbit (1:400; Vector Biolabs BA-1000), AlexaFluor 647 anti-mouse (1:200; Invitrogen A21235), and AlexaFluor 568 anti-rat (1:400; Invitrogen A11077). The GFP signal was amplified with extra steps of incubation with ABC Elite kit (1:200; Vector Laboratories) and TSA^TM^ Fluorescein Tyramide Reagent kit (1:50; AKOYA Biosciences SAT701001EA). For BrdU staining, sections were denatured in 2N HCL at 30°C for 30 minutes. The 2- and 7-DPI samples were denatured before primary antibody incubation. The 4- and 8-WPI samples were stained for GFP and NeuN first, followed by denaturing and BrdU staining (**Figure 1c**)

### 2.5. Stereological Cell Quantification and Statistics

Stereology was used for total cell estimation for all groups using **N = ΣQ^−^.t/h ·1/asf·1/ssf**, where **Q^−^** is cells counted, **t** is measured section thickness, **h** is counting frame height, **asf** is area sampling fraction, and **ssf** is section sampling fraction (Zhao & van Praag, 2020). For cell proliferation study of 2- and 7-DPI groups, an inverted light microscope (1X71 Olympus) was used to view samples and quantify the number of BrdU+ cells. The Visiopharm (Olympus) program was used to count total number of BrdU+ cell in the granular cell layers and hilus as we previously described (Sun et al. 2007). For 4- and 8-WPI study groups, an LSM710 confocal microscope was used to capture images of the entire DG at 20x magnification. The Z-stack and tiling functions in Zeiss Zen microscope software was used to capture a stack made up of 10 images, each spanning a total of 1µm and all adjacent to each other for a total stack size of 10µm. Z-stack files were put into FIJI and merged into a Z-projection. Spectral unmixing was conducted on each image to remove the bleed-through of channels that were used to capture fluorescent labeling. The total number of cells were counted in the GCLs region using the FIJI plugin Cell Counter.

For data analysis, stereologically calculated cell numbers for all data were evaluated using two-way ANOVA (JMP Pro v16) combined with Tukey HSD post hoc tests. For group comparison, the Grubb’s Outlier test was used and the outliers were removed from the dataset. The significance level was set to α = 0.05 for all analyses performed, and averaged values are expressed as mean ± SEM.

## 3. Results

### 3.1. Notch1 deletion in Nestin+ cells abolished the TBI-enhanced cell proliferative response in the neurogenic region of the DG at the acute stage following injury

In this study, we first assessed the effect of TBI on cell proliferative responses in the DG at the acute (2 DPI) and subacute (7 DPI) stage and Notch1 deficiency in NSCs affecting this response. Proliferative cells were labeled with pulse BrdU injection, and the number of BrdU+ cells in the hippocampal DG granular cell layer (GCL, including the subgradular zone) and hilus on the hemisphere ipsilateral to the injury was quantified using the unbiased stereological quantification method (**Figure 2a**).

**Figure 2.**
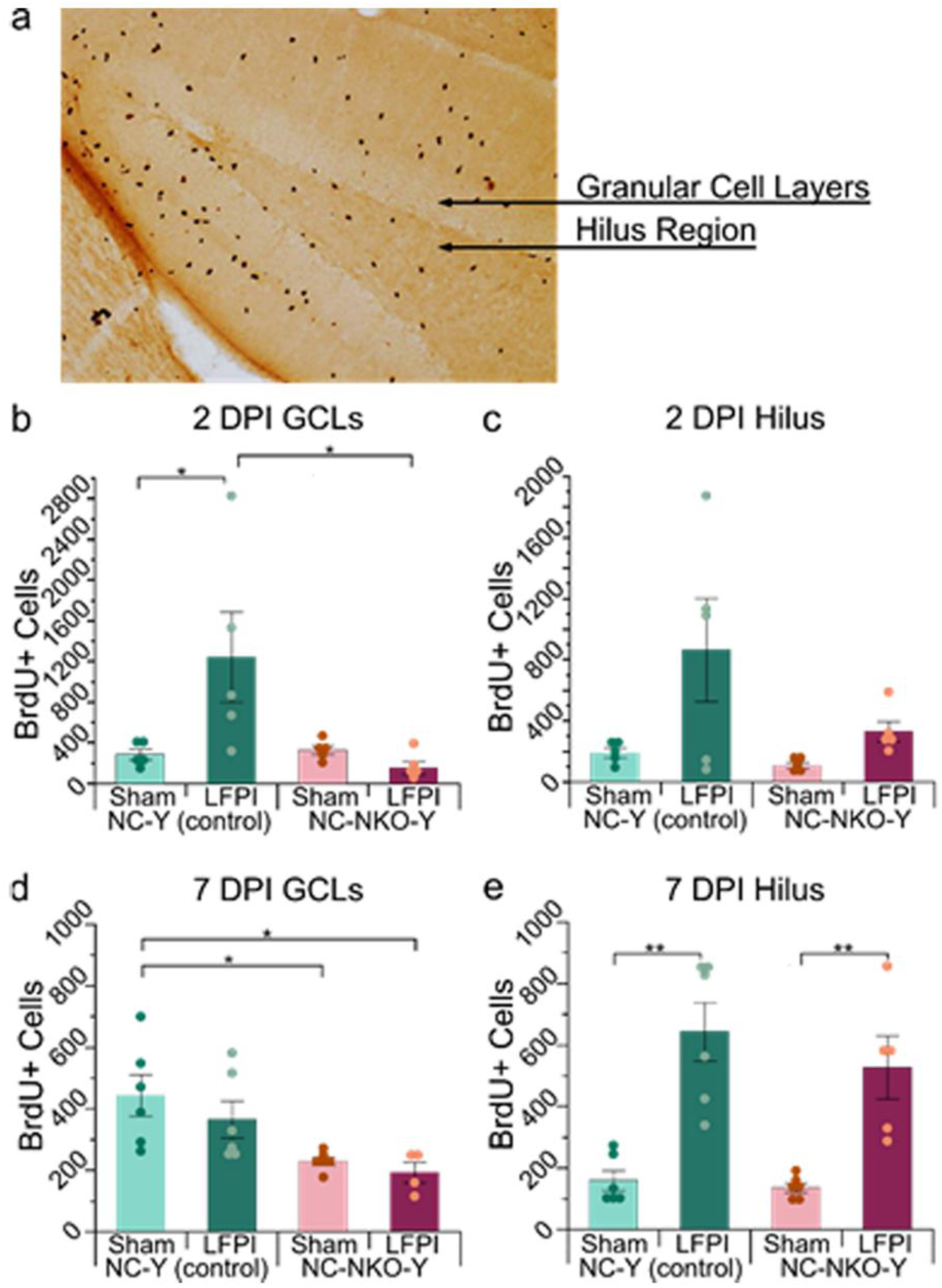
Injury induces cell proliferation in the GCL of the DG at the acute stage following TBI and this Notch1 deletion blockes this response. (**a**). The representative image of BrdU stained (brown dots) coronal section showing the two regions analyzed, the GCLs and hilus regions of the DG. Group means for stereologically calculated total number of BrdU+ cell are plotted for (**b**) 2 DPI GCLs, (**c**) 2 DPI hilus, (**d**) 7 DPI GCLs, and (**e**) 7 DPI hilus. Significance levels indicated by \**p* < 0.05, \*\**p* < 0.005.

Two-way ANOVA Tukey HSD post hoc test has shown that at the acute stage following TBI (2 DPI, **Figure 2b&c**), in the control mice (NC-Y line), a significant increase in the number of BrdU+ cells was observed in the GCL in the injured group compared to the matched sham (**Figure 2b**; *p* = 0.0168*). In the Notch1 cKO mice (NC-NKO-Y line), TBI did not increase the number of BrdU+ cells in either the GCL or hilus. When comparing by genotype between the control and Notch1 lines, no difference was found in sham groups, however, a significantly lower number of BrdU+ cells was observed in the GCL in the injured Notch1 cKO mice when compared to the control injured group (**Figure 2b**, p = 0.01*). At this time point, the number of BrdU+ cells in the hilus region has no significant difference related to injury or genotype.

At the subacute stage following TBI (7 DPI, **Figure 2d&e**), in the control mice (NC-Y line), a significantly higher number of BrdU+ cells was found in the hilus region in the injured group compared to the matched sham (**Figure 2d**, *p* < 0.0001**), whereas no difference was found in the GCL. In the Notch1 cKO mice, a similar pattern as the control mice was found with the injured group having higher number of BrdU+ cells in the hilus region not the CGL than the matched sham group (**Figure 2e**, *p* = 0.0094**). When comparing by genotype, a significantly lower number of BrdU+ cells was found in the Notch1 cKO mice in the GCL in both the sham and injured groups compared to the sham control mice (**Figure 2d**, sham NC-Y vs. sham NC-NKO-Y, *p* = 0.0304*; sham NC-Y vs. LFPI NC-NKO-Y, *p* = 0.0212*).

Collectively, the BrdU pulse labeling data revealed that moderate LFPI enhanced cell proliferation in the GCL of the DG at the acute stage following TBI and this injury-induced cell proliferative response was abolished with Notch1 conditionally deleted from the nestin+ NSCs. At the subacute stage following TBI, injury-enhanced cell proliferation was more prominent in the hilus region and this was not affected by Notch1 deletion from the nestin+ NSCs, however, Notch1 cKO significantly affected the number of BrdU-labeled proliferative cells in the GCL.

### 3.2. Survival and neuronal differentiation of injury-induced new cells in the DG was not affected by Notch1 deletion

Long-term survival and neuronal differentiation of injury-induced new cells are critical for these cells to contribute to post-injury brain repair and regeneration. To evaluate the influence of TBI and Notch1 deletion from NSCs on survival and neuronal differentiation of injury-induced new cells in the DG, we counted the number of BrdU+ cells and BrdU/NeuN double-labeled cells that were generated at 1–7 DPI and survived to 4- or 8-WPI in the GCL. As new neurons generated in the DG migrate gradually from the subgranular zone laterally to the outer layer of the GCL, we divided the GCL into three zones (inner, middle, and outer thirds extending from the subgranular zone designated as 1, 2, and 3 respectively) following a published principle (Kempermann et al. 2003), for counting BrdU+ and BrdU+/NeuN+ cells to assess the influence of injury on cell migration.

At 4 WPI, the time point when injury-induced new cells starting to express mature neuronal marker NeuN, a significant overall injury effect was observed with higher number of BrdU+ [F(1,16) = 11.08; *p* = 0.0043**] and BrdU+/NeuN+ [F(1,16) = 9.638; *p* = 0.0068**] as a result of injury (**Figure 3a,c; supplemental data-Table 1**). This injury effect was observed in all three GCL zones (**supplemental data-Table 2**). At this time point, the difference due to genotype effect was not observed for BrdU+ [F(1,16) = 0.133; *p* = 0.721], nor BrdU+/NeuN+ [F(1,16) = 0.147; *p* = 0.706], and there was no interaction between injury and genotype for BrdU+ [F(1,16) = 0.216; *p* = 0.648] and BrdU+/NeuN+ [F(1,16) = 0.195; *p* = 0.665]. Although the number of BrdU+, and BrdU+/NeuN+ cells were higher in the injured groups, due to variables, the post-hoc test did not reveal significant difference between sham and injured groups within the same mice lines.

**Figure 3.**
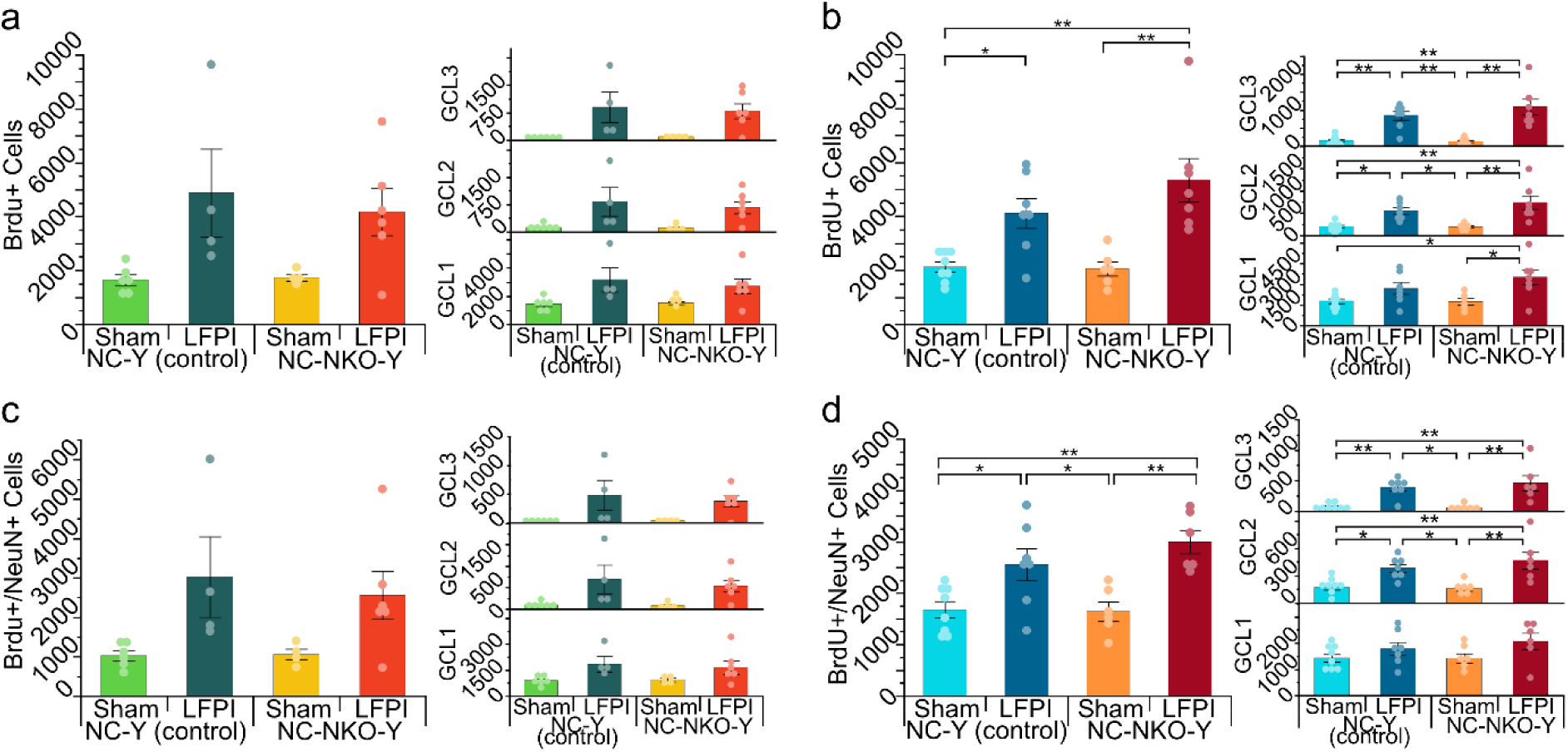
Survival of injury-induced new cells at 4 and 8 WPI. (**a-b**). Group means for stereologically calculated total cell number of BrdU+ cells at 4 WPI (**a**) and 8 WPI (**b**) in entire GCL and sub-regions of GCL. (**c-d**) Total number of BrdU+/NeuN+ cells at 4 WPI (**c**) and 8 WPI (**d**) in entire GCL and sub-regions of GCL. Injured animals had significantly higher number of BrdU+ and BrdU+/NeuN+ cells. Significance levels indicated by \**p* < 0.05, \*\**p* < 0.005.

At 8 WPI, the time point when new neurons in the DG generated following TBI become mature neurons (Sun et al. 2007), a robust overall injury effect was observed with higher number of both BrdU+ [F(1,24) = 26.358; *p* < 0.0001**] and BrdU+/NeuN+ [F(1,23) = 23.759; *p* < 0.0001*] as a result of injury (**Figure3 b,d; supplemental data-Table 1 and 3**). Post-hoc analysis also revealed the injury effect in both mice lines. For BrdU+ cells, injured groups had significantly higher number than matched sham groups (LFPI NC-Y vs. sham NC-Y, *p* = 0.0421*; LFPI NC-NKO-Y vs. sham NC-NKO-Y, p = 0.0011**). A similar injury effect was also observed for BrdU+/NeuN+ cells, with injured groups having a significantly higher number of cells than the matched sham groups (LFPI NC-Y vs. sham NC-Y, *p* = 0.0396*; LFPI NC-NKO-Y vs. sham NC-NKO-Y, *p* = 0.0032**). When BrdU+ and BrdU+/NeuN+ cells were divided into three GCL zones, a significantly higher number of BrdU+ and BrdU+/NeuN+ cells were observed in the middle and outer zones of GCL (GCL2 and 3) in the injured groups compared to matched sham in both mice lines. Similar to 4 weeks after injury, genotype did not affect the amount of surviving BrdU+ [F(1,24) = 1.289; *p* = 0.268] or BrdU+/NeuN+ [F(1,23) = 0.804; *p* = 0.379] cells, nor were there interactions for BrdU+ [F(1,24) = 1.605; *p* = 0.217] or BrdU+/NeuN+ [F(1,23) = 1.069; *p* = 0.312].

### 3.3. Injury affected fate choice of nestin+ NSCs in the DG at the chronic stage following TBI

The mouse lines used in this study have a fluorescent reporter for nestin+ NSCs including the conditional deletion of Notch1 from these cells within the Notch1 cKO line. In both the control and Notch1 cKO lines, cells with GFP+ labeling originated from the nestin+ population tagged with the eYFP fluorescent reporter. To assess the influence of injury on the nestin+ NSC pool where the neurogenic response originated, we counted the number of total GFP+ cells in the DG in a defined counting frame in animals survived for 4 or 8 weeks post-injury. These assessments were made on the number of all GFP+ cells (**Figure 4**), regardless of phenotypic expression of the other co-labeling factors.

**Figure 4.**
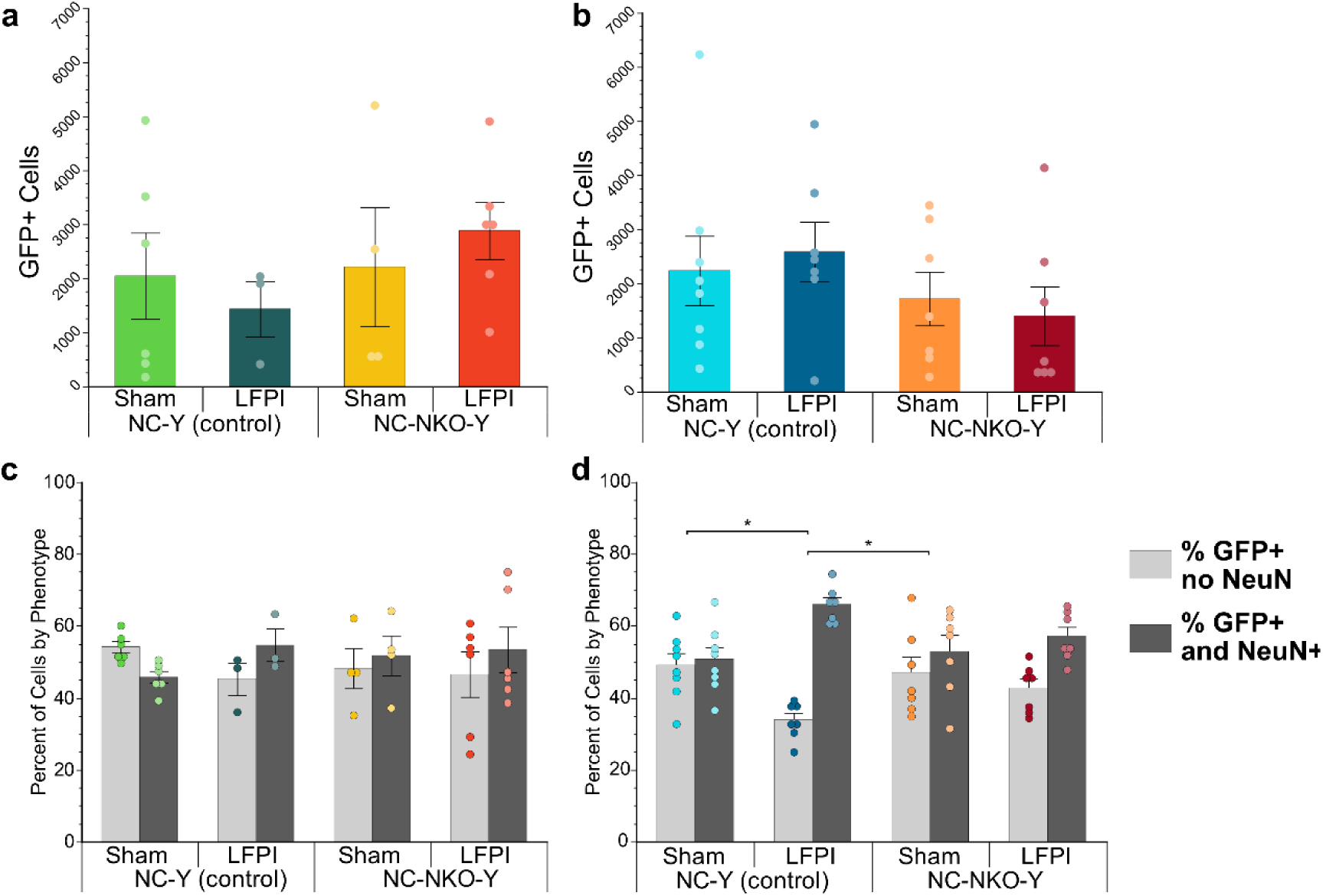
Neuronal differentiation from GFP+ populations is increased following TBI at 8 WPI. (**a-b**). Means for stereologically calculated total number of cell are plotted for GFP+ cells at 4 WPI (**a**) and 8 WPI (**b**) in entire GCLs. No significant differences were observed at both time points. (**c-d**). The percentage of GFP+/NeuN-cells and GFP+/NeuN+ cells for both 4 WPI (**c**) and 8 WPI (**d**) in all GCLs. Significant differences were observed at 8WPI between the injured NC-Y group compared to both sham NC-Y and sham NC-NKO-Y groups (\**p* < 0.05).

At 4 or 8 WPI, the total number of GFP+ cells (**Figure 4a**) did not show difference as a result of injury [F(1,15) = 0.0014; *p* = 0.971 for 4 WPI; [F(1,25) = 0.00089; *p* = 0.978] for 8WPI], or genotype [F(1,15) = 0.976; *p* = 0.339for 4WPI; [F(1,25) = 0.343; *p* = 0.564]] in the GCL as a whole.

In addition to the total number of GFP+ cells, we differentially quantified the number of GFP+ cells that were NeuN+ as this elucidated the fate choice of the nestin+ NSC populations to neuronal differentiation. At 4 WPI (**Figure 4c**), no difference was found by injury [F(1,15) = 1.054; *p* = 0.321], or genotype [F(1,15) = 0.217; *p* = 0.648]. However, at 8 WPI (**Figure 4d**), injured control mice (NC-Y line) had a significantly higher proportion of GFP+/NeuN+ cells as a result of injury [F(1,25) = 9.231; *p* = 0.005*], with 66% of GFP+ cells expressing NeuN+ cells compared to 50% in sham control, 52% in sham Notch1 cKO mice and 57% in injured Notch1 cKO mice (NC-NKO-Y line) (**supplemental data Table 5**). This difference in proportions was not significant by genotype comparisons [F(1,25 = 1.097; *p* = 0.305], or interactions [F(1,25) = 2.939; *p* = 0.0988]. This result shows that injury affects the fate choice of nestin+ NSCs with higher percentage of cells committing neuronal differentiation.

### 3.4. TBI or Notch1 deletion drove injury-induced dividing nestin+ NSCs in the DG towards neuronal differentiation

In the neurogenic region of the DG, the BrdU-labeled dividing cells included both fast diving progenitor cells and slow dividing NSCs. Nestin is the most widely used marker for NSCs. In our mouse lines, nestin+ cells were tagged with GFP, thus BrdU+/GFP+ cells representing nestin+ NSCs undergoing division at 1-7 DPI. To assess the influence of TBI and Notch1 cKO on the survival of this cell population, we quantified the number of BrdU+/GFP+ cells at 4 and 8 weeks post-TBI. Our stereological quantification revealed that at 4 WPI, the number of BrdU+/GFP+ in the GCL did not show a difference as a result of injury [F(1,16) = 1.176; *p* = 0.294] or by genotype [F(1,16) = 1.026; *p* = 0.326] (**Figure 5a**). Similarly, at 8 WPI, no difference was found in the number of BrdU+/GFP+ in the GCL by injury [F(1,25) = 0.0005; *p* = 0.982] or by genotype [F(1,25) = 0.123; *p* = 0.729] (**Figure 5b**). This suggests that injury or Notch1 cKO did not affect the number of nestin+ NSCs that were dividing at 1-7 DPI and survived for 4- or 8-WPI.

**Figure 5.**
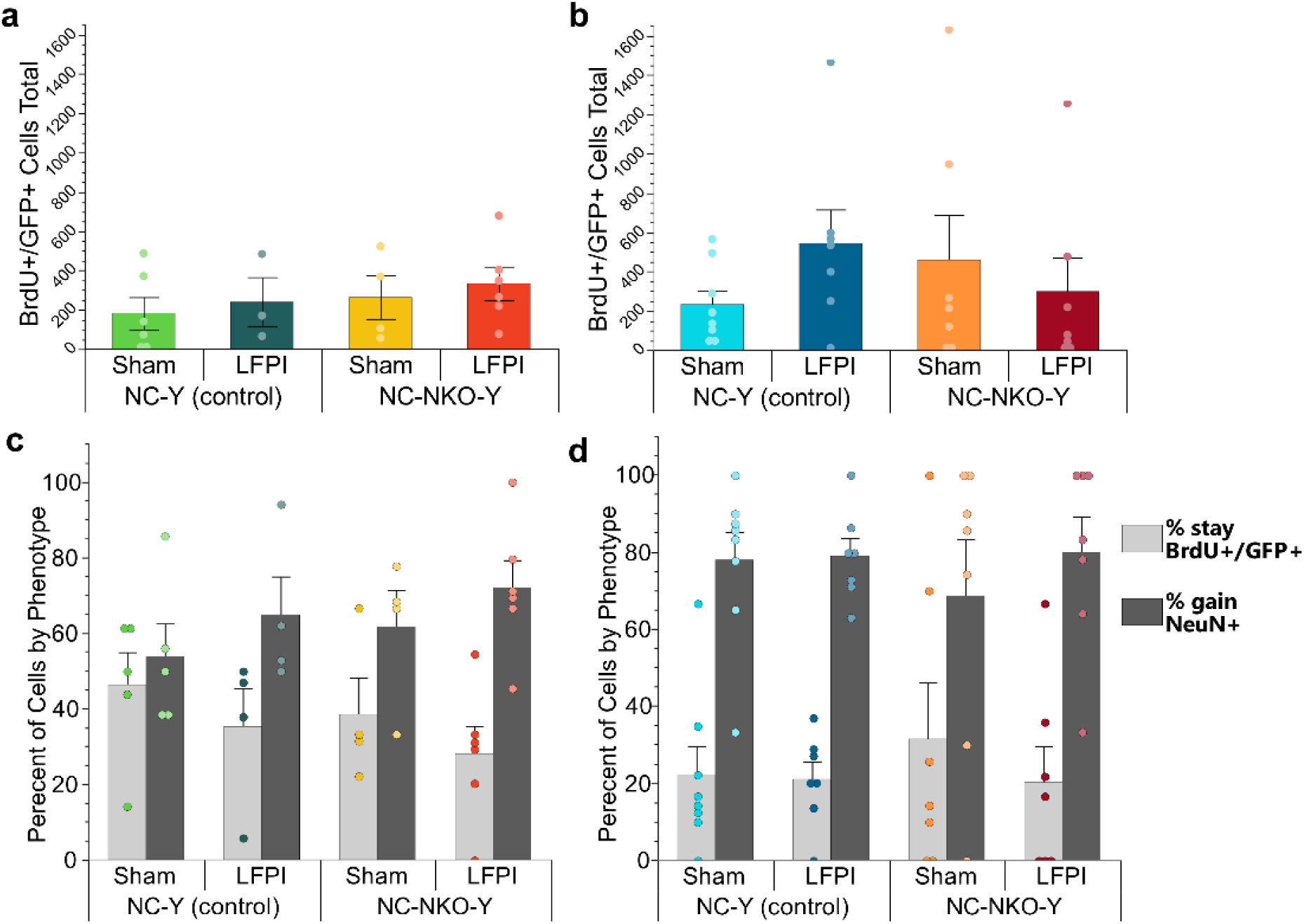
Loss of Notch1 does not alter injury-induced neuronal populations. Group means for stereologically calculated cell amounts are plotted for all BrdU+/GFP+ cells, including both NeuN- and NeuN+, at (**a**) 4 WPI and (**b**) 8 WPI. Total percentage of cells by phenotype with BrdU+/GFP+/NeuN-as the left-hand bars in light gray and BrdU+/GFP+/NeuN+ as the right-hand bars in dark gray for both (**c**) 4 WPI and (**d**) 8 WPI in all GCLs. No differences were observed between the separate conditions.

To assess the influence of TBI and Notch1 cKO on neuronal fate choice of nestin+ NSCs that were driven into S1 phase at 1-7 DPI, we also qualified the percentage of these cells becoming mature neurons (BrdU+/GFP+/NeuN+ cells). At 4 WPI, the percentage of BrdU+/GFP+/NeuN-cells and BrdU+/GFP+/NeuN+ cells were at 46.5% vs 53.5% in NC-Y sham group, 35% vs 65% in NC-Y injured group, 38% vs 62% in NC-NKO-Y sham group and 28% vs 72% in NC-NKO-Y injured group (**Figure 5c**; **supplemental data Table 5**). A shifting of higher percentage of BrdU/GFP+ cells becoming NeuN+ was noticed in relation to injury and Notch 1 cKO. At 8 WPI, the majority of BrdU+/GFP+ cells became mature neurons triple-labeled with NeuN (BrdU+/GFP+/NeuN+) in all four study groups (**Figure 5d, supplemental data Table 5**). These data suggest that TBI and Notch1 cKO potentially drives dividing NSCs towards neuronal fate.

## 4. Discussion

Neurogenesis is widely recognized as the brain’s endogenous reparative response to neuropathological conditions including TBI. The extent of this injury-induced cellular response particularly in the hippocampal region has been examined in different injury models at different injury intensities due to their contribution to learning and memory function (Blaiss et al. 2011; Chirumamilla et al. 2002; Sun et al. 2007; Sun et al. 2015). Most of the published studies examining the impact of TBI on hippocampal neurogenesis have used thymidine analogs BrdU or edU to trace injury-induced cell activation for a limited period (Chirumamilla et al. 2002; Rice et al. 2003; Dash, Mach, and Moore 2001; Sun et al. 2009). Neurogenesis is a dynamic response involving both slow dividing NSCs and fast dividing progenitor cells, as thymidine analogs can only tag dividing cells at a short period following administration, thus providing a limited view of injury-induced changes on this response. In the current study, we used transgenic animals that have induced eYFP expression in nestin+ NSCs in combination with BrdU labeling to precisely track the injury effect on nestin+ NSCs for a prolonged period. This study also examined the role of Notch1 in regulating nestin+ NSC proliferation, survival and differentiation following TBI. Using these unique transgenic mice lines, we examined the injury-induced nestin+ NSC response at the acute, subacute and chronic stage following TBI. We found that moderate LFPI enhanced cell proliferation in the DG at the acute stage following LFPI in mice. However, this injury-induced response was abolished when Notch 1 was conditionally deleted from nestin+ NSCs. At the chronic stage following TBI, injury-induced DG new cells persisted at higher level with a majority of cells becoming mature neurons, while the survival and neuronal differentiation of injury-induced new cells were not affected by Notch1 deletion. Additionally, both TBI and Notch1 deletion drove NSC into neuronal differentiation. These data confirm the impact of TBI on hippocampal neurogenesis post-injury and the role of Notch1 in regulating this cellular response in the injured brain.

The injury effect on NSC proliferation, survival and neuronal differentiation in the control mouse line is anticipated as the general trend and is in agreement with what we previously reported in the rat LFPI model (Sun et al. 2005; Sun et al. 2007). A slight difference in the cell proliferative response is observed at the subacute stage of 7 DPI (**Figure 2d&e**). In our rat LFPI model, injury-enhanced cell proliferation is observed at both 2- and 7-DPI with the peak time at 2 days (Sun et al. 2005). In the current mouse LFPI model, injury-enhanced cell proliferation is only observed at 2 but not 7 DPI in the GCL where the NSCs reside. At 7 DPI, a higher number of BrdU+ cells are predominantly located in the hilus region. This difference maybe due to species differences and variable injury levels, as injury severity has a significant impact on the NSC response (Gao, Enikolopov, and Chen 2009; Wang et al. 2016). This study has also demonstrated that Notch1 deletion from nestin+ NSCs only affects injury-enhanced NSC proliferation in the GCL region, whereas injury-induced glial cell proliferation in the hilus region is not affected. The Notch1 signaling pathway regulates proliferation and differentiation of NSCs in the developmental and adult brain (Ables et al. 2010; Androutsellis-Theotokis et al. 2006; Imayoshi et al. 2010; Zhang et al. 2015); however, little is known about its role in regulating adult neurogenesis post-TBI. Our finding that loss of Notch1 in nestin+ NSCs leads to a reduction of injury-induced cell proliferation in the DG supports the notion that Notch1 is an essential player dictating NSC proliferation, not only under physiological conditions, but also under neuropathological condition.

For injury-induced new cells participating regeneration, cells need to survive long-term and become mature functional neurons. It is known that under physiological conditions newly generated granule cells in the DG have specific developmental milestones that can serve as checkpoints for survival, and a large percentage of cells undergo apoptosis at 2-4 weeks after birth (Beining et al. 2017; Cole et al. 2020; Zhao, Song, et al. 2006). Cells that survived this selection process express the mature neuronal marker NeuN around 4 weeks and become indistinguishable from neighboring pre-existing mature granule neurons around 8 weeks after birth (Zhao, Teng, et al. 2006). In our study, we have tracked the survival and neuronal differentiation of injury-induced proliferative nestin+ NSCs in the DG at 4- and 8-WPI by counting the number of BrdU+ and BrdU+/NeuN+ cells (**Figure 3**). We have found that a higher number of BrdU+ alone or BrdU/NeuN double-labeled cells persist in the injured control mouse line and predominantly in the middle and outer zones of the GCL, confirming the long time survival, neuronal differentiation and lateral migration of injury-induced proliferating cells. This observation is in agreement with our previous report in the rat LFPI model (Sun et al. 2007). What was unexpected is that a higher number of BrdU+ cells is also observed in the injured Notch1 cKO mouse line. In the proliferation study group, Notch 1 cKO mice have a reduced number of BrdU+ cells in the CGL at 2- and 7-DPI, we therefore would expect a lower number of BrdU+ cells at the chronic stage as well. This discrepancy is probably due to the following reasons: 1) different BrdU injection paradigm. The animals at 2- and 7-DPI groups received single pulse BrdU injection at 2 hours before being sacrificed, whereas the animals survived to the chronic stage received continuous BrdU injection at 1-7 DPI, thus a lot more proliferating cells were BrdU labeled and survived. 2). Notch signaling functions to maintain NSCs pool by instructing NSC asymmetric division where one daughter remains as a NSC and the other one going on a differentiation fate. Loss of Notch results in both daughter cells of dividing NSCs committing to neuronal differentiation thus higher number of BrdU+ and BrdU+/NeuN+ cells at the chronic stage following injury (Castro et al. 2006). Our data suggest that Notch1 deletion in nestin+ NSCs does not affect survival of injury-induced new cells in the DG.

In this study, we have also examined the number of GFP+ cell, and the ratio of GFP+ cells co-labeled with NeuN+ at 4- and 8-WPI. In our study, nestin+ NSCs are tagged with GFP and these cells have lost Notch1 expression upon TAM treatment in the Notch 1 cKO mouse line. When counting the total GFP+ cell number in our counting frame, no significant difference was found between injured and sham animals in both control and Notch cKO lines in both 4 and 8 WPI (**Figure 4 a&b**). When examining the ratio of GFP+ cells becoming mature neurons at these two time points (GFP+/NeuN+ cells), we found a higher percentage of neuronal differentiation at 8 WPI only in the injured control mice. This result suggests that injury affects neuronal fate choice of nestin+ NSCs, which is in agreement to our previous published study in a rat LFPI model (Sun et al. 2007).

In the normal neurogenic process in the DG, dividing nestin+ NSCs differentiate into neurons and astrocytes proportionally (Filippov et al. 2003; Fukuda et al. 2003; Artegiani et al. 2017). When examining neuronal fate choice of GFP+ NSCs that underwent division (GFP+/BrdU+ cells) and those not (GFP+/BrdU-cells) at 1-7 DPI, we found that injury and/or Notch1 deletion induces GFP+/BrdU+ NSCs committing an earlier neuronal differentiation at 4 WPI (GFP+/BrdU+/NeuN+ cells, **Figure 5d**), whereas injury alone drives the overall nestin+ NSCs into neuronal differentiation at more chronic stage following TBI at 8 WPI (GFP+/NeuN+/BrdU-cells, **Figure 5**). These data suggest that both TBI and Notch1 cKO push nestin+ NSCs towards neuronal fate. The injury effect on nestin+ NSC fate choice towards neuronal phenotype has not directly been reported before. A recently published transcriptomic study has reported that at 15 DPI, a focal brain injury (cortical impact injury) affects fate commitment of NSCs in the DG by promoting neurogenesis while decreasing astrogliogenesis at the transcriptional level (Bielefeld et al. 2024). Our data is in agreement with this study though at the cellular level. The influence of Notch1 deletion on inducing nestin+ NSC neuronal differentiation is in agreement with the established role of Notch1 functioning on NSCs fate choice. In the normal neurogenic response, Notch signaling activation maintains NSCs in a balanced state of self-renewal, proliferation and differentiation via regulating expression of transcription factors of Hes1 and Hes5 to repress the expression of proneural genes (Artavanis-Tsakonas, Rand, and Lake 1999; Kageyama et al. 2008; Ohtsuka et al. 1999). Inhibition of Notch signaling is known to down-regulate Hes1 and up-regulate proneural genes inducing neuronal differentiation (Kageyama et al. 2008; Shimojo, Ohtsuka, and Kageyama 2008). Under neuropathological conditions, the role of Notch 1 in regulating NSC fate choice is less known. In a spinal cord injury model, genetic knockdown of Notch1 signaling or pharmacologically blocking Notch1 signaling can induce endogenous reactive astrocytes to reprogram into neurons in the injured spinal cord in adult mice (Tan et al. 2022). In a focal brain injury model, downregulation of Hes1 via RNA interference promotes the differentiation of neural progenitor cells into mature neurons in the DG (Zhang et al. 2014). Similar to these published studies, our study with conditional deletion of Notch1 from nestin+ NSCs drives proliferating NSCs into neuronal differentiation in both sham and injured Notch1 cKO mice.

## 5. Conclusion

Adult hippocampal neurogenesis plays important roles in hippocampal-dependent learning and memory functions (Eriksson et al. 1998; Anacker and Hen 2017; Aimone, Deng, and Gage 2011; Deng, Aimone, and Gage 2010; Marin-Burgin and Schinder 2012). TBI induces a heightened level of hippocampal neurogenesis, and this response is believed to aid post-injury cognitive recovery (Chirumamilla et al. 2002; Blaiss et al. 2011; Sun et al. 2007; Sun et al. 2015). It is hoped that this endogenous cellular response can be stimulated to promote brain repair and regeneration. To achieve this, it is necessary to fully understand the extent of the injury-induced neurogenic response and regulating mechanisms. The current study examined the influence of TBI on nestin+ NSC proliferation, survival and neuronal differentiation in the injured hippocampus, and the role of Notch1 in regulating these processes. We have found that a moderate TBI augments proliferation of NSCs at the acute stage following TBI and induces these cells to become mature neurons, and Notch1 is involved in regulating injury-induced NSC proliferation and neural differentiation.

## CRediT Authorship contribution statement

Nicole Weston: investigation, methodology, visualization, data acquisition and analysis, writing original draft Timothy N. Keoprasert: investigation, methodology Jakob Green: investigation, methodology Sarah Baig: investigation, methodology Dong Sun: conceptualization, supervision, writing – review, editing and final draft, funding acquisition, project administration

## Ethics approval statement

All animal procedures were conducted following the “Principles of laboratory animal care” (NIH publication No. 86-23, revised 1985)

## Data availability

Data are available from the corresponding author on reasonable request.

## Supplemental Data

**Table 1.** Significance Values for Comparisons of 4- and 8-week post-Injury. The cell populations of the different variations in phenotypic expressions (left column) compared by injury, genotype, or an interaction of injury and genotype with *p* < 0.005 designated in orange. Data is separated by 4 WPI (top) and 8 WPI (bottom).

| Cell Phenotype | Injury | Genotype | Injury*Genotype |
| --- | --- | --- | --- |
| **4 WPI** |  |  |  |
| **BrdU+** | $p = 0.0043^{**}$ | $p = 0.721$ | $p = 0.648$ |
| **GFP+** | $p = 0.971$ | $p = 0.339$ | $p = 0.443$ |
| **BrdU+/GFP+** | $p = 0.294$ | $p = 0.326$ | $p = 0.254$ |
| **BrdU+/NeuN+** | $p = 0.0068^{**}$ | $p = 0.706$ | $p = 0.665$ |
| **GFP+/NeuN+** | $p = 0.317$ | $p = 0.947$ | $p = 0.799$ |
| **BrdU+/GFP+/NeuN+** | $p = 0.111$ | $p = 0.495$ | $p = 0.281$ |
| **8 WPI** |  |  |  |
| **BrdU+** | $p < 0.0001^{**}$ | $p = 0.268$ | $p = 0.217$ |
| **GFP+** | $p = 0.978$ | $p = 0.146$ | $p = 0.563$ |
| **BrdU+/GFP+** | $p = 0.982$ | $p = 0.729$ | $p = 0.129$ |
| **BrdU+/NeuN+** | $p < 0.0001^{**}$ | $p = 0.379$ | $p = 0.312$ |
| **GFP+/NeuN+** | $p = 0.257$ | $p = 0.072$ | $p = 0.137$ |
| **BrdU+/GFP+/NeuN+** | $p = 0.283$ | $p = 0.087$ | $p = 0.085$ |

**Table 2.** Significance Values for Comparisons of 4 WPI Separated by GCL. The cell populations of the different variations in phenotypic expressions (left column) compared by injury, genotype, or an interaction of injury and genotype with *p* < 0.05 designated in red and *p* < 0.005 designated in orange. Data is separated by GCL 1 (top), GCL 2 (middle), GCL 3 (bottom).

| Cell Phenotype | Injury | Genotype | Injury*Genotype |
| --- | --- | --- | --- |
| **4 WPI** |  |  |  |
| **GCL 1** |  |  |  |
| **BrdU+** | **$p = 0.0091^{**}$** | $p = 0.702$ | $p = 0.591$ |
| **GFP+** | $p = 0.911$ | $p = 0.347$ | $p = 0.578$ |
| **BrdU+/GFP+** | $p = 0.320$ | $p = 0.325$ | $p = 0.257$ |
| **BrdU+/NeuN+** | **$p = 0.0171^{*}$** | $p = 0.807$ | $p = 0.699$ |
| **GFP+/NeuN+** | $p = 0.125$ | $p = 0.416$ | $p = 0.236$ |
| **BrdU+/GFP+/NeuN+** | $p = 0.125$ | $p = 0.416$ | $p = 0.236$ |
| **GCL 2** |  |  |  |
| **BrdU+** | **$p = 0.0038^{**}$** | $p = 0.644$ | $p = 0.697$ |
| **GFP+** | $p = 0.379$ | $p = 0.414$ | $p = 0.180$ |
| **BrdU+/GFP+** | $p = 0.157$ | $p = 0.529$ | $p = 0.330$ |
| **BrdU+/NeuN+** | **$p = 0.0041^{**}$** | $p = 0.592$ | $p = 0.666$ |
| **GFP+/NeuN+** | $p = 0.091$ | $p = 0.663$ | $p = 0.529$ |
| **BrdU+/GFP+/NeuN+** | $p = 0.091$ | $p = 0.663$ | $p = 0.530$ |
| **GCL 3** |  |  |  |
| **BrdU+** | **$p = 0.0022^{**}$** | $p = 0.844$ | $p = 0.764$ |
| **GFP+** | $p = 0.483$ | $p = 0.546$ | $p = 0.197$ |
| **BrdU+/GFP+** | $p = 0.224$ | $p = 0.224$ | $p = 0.224$ |
| **BrdU+/NeuN+** | **$p = 0.0048^{**}$** | $p = 0.674$ | $p = 0.653$ |
| **GFP+/NeuN+** | $p = 0.231$ | $p = 0.272$ | $p = 0.231$ |
| **BrdU+/GFP+/NeuN+** | $p = 0.220$ | $p = 0.272$ | $p = 0.231$ |

**Table 3.** Significance Values for Comparisons of 8 WPI Separated by GCL. The cell populations of the different variations in phenotypic expressions (left column) compared by injury, genotype, or an interaction of injury and genotype with *p* < 0.05 designated in red and *p* < 0.005 designated in orange. Data is separated by GCL 1 (top), GCL 2 (middle), GCL 3 (bottom).

| Cell Phenotype | Injury | Genotype | Injury*Genotype |
| --- | --- | --- | --- |
| **8 WPI** |  |  |  |
| **GCL 1** |  |  |  |
| **BrdU+** | $p = 0.0015^{**}$ | $p = 0.317$ | $p = 0.268$ |
| **GFP+** | $p = 0.739$ | $p = 0.145$ | $p = 0.639$ |
| **BrdU+/GFP+** | $p = 0.616$ | $p = 0.886$ | $p = 0.163$ |
| **BrdU+/NeuN+** | $p = 0.0365^{*}$ | $p = 0.541$ | $p = 0.520$ |
| **GFP+/NeuN+** | $p = 0.595$ | $p = 0.081$ | $p = 0.166$ |
| **BrdU+/GFP+/NeuN+** | $p = 0.611$ | $p = 0.066$ | $p = 0.079$ |
| **GCL 2** |  |  |  |
| **BrdU+** | $p < 0.0001^{**}$ | $p = 0.332$ | $p = 0.299$ |
| **GFP+** | $p = 0.0321^{*}$ | $p = 0.261$ | $p = 0.361$ |
| **BrdU+/GFP+** | $p = 0.494$ | $p = 0.356$ | $p = 0.278$ |
| **BrdU+/NeuN+** | $p < 0.0001^{**}$ | $p = 0.497$ | $p = 0.364$ |
| **GFP+/NeuN+** | $p = 0.0051^{**}$ | $p = 0.157$ | $p = 0.105$ |
| **BrdU+/GFP+/NeuN+** | $p = 0.135$ | $p = 0.559$ | $p = 0.413$ |
| **GCL 3** |  |  |  |
| **BrdU+** | $p < 0.0001^{**}$ | $p = 0.391$ | $p = 0.316$ |
| **GFP+** | $p = 0.142$ | $p = 0.837$ | $p = 0.365$ |
| **BrdU+/GFP+** | $p = 0.310$ | $p = 0.310$ | $p = 0.310$ |
| **BrdU+/NeuN+** | $p < 0.0001^{**}$ | $p = 0.671$ | $p = 0.536$ |
| **GFP+/NeuN+** | $p = 0.0796$ | $p = 0.594$ | $p = 0.996$ |
| **BrdU+/GFP+/NeuN+** | $p = 0.115$ | $p = 0.899$ | $p = 0.826$ |

**Table 4.** Summary of marker expression of NSCs at 4- and 8-WPI. Separate cell phenotype combinations (left column) with a generalized summary of results (right column) for 4 WPI (top) and 8 WPI (bottom). Arrows indicate an increase or decrease.

| Cell Phenotype | Result |
| --- | --- |
| **4 WPI** |  |
| BrdU+ | ↑ from injury |
| GFP+ | - |
| BrdU+ / GFP+ | - |
| BrdU+ / NeuN+ | ↑ from injury |
| GFP+ / NeuN+ | - |
| **8 WPI** |  |
| BrdU+ | ↑ from injury |
| GFP+ | ↓ from injury |
| BrdU+ / GFP+ | - |
| BrdU+ / NeuN+ | ↑ from injury |
| GFP+ / NeuN+ | trend ↓ from Notch1 cKO |

**Table 5.** Percentage of NeuN Expressing Cell Populations of 4- and 8-week post-Injury. The cell populations of the different variations in phenotypic expression (left column) compared by genotype and injury condition. Data is separated by proportions of the total GFP+ population expressing NeuN and proportions of the injury-induced NSC population (BrdU+/GFP+) expressing NeuN.

| Cell Phenotype | NC-Y (control)<br>Sham | NC-Y (control)<br>LFPI | NC-NKO-Y<br>Sham | NC-NKO-Y<br>LFPI |
| --- | --- | --- | --- | --- |
| **4 WPI** |  |  |  |  |
| **GFP+ Population: NeuN-** | 54.25% | 45.30% | 48.20% | 46.53% |
| **GFP+ Population: NeuN+** | 45.75% | 54.70% | 51.80% | 53.47% |
| **BrdU+/GFP+/NeuN-** | 46.50% | 35.22% | 38.45% | 28.03% |
| **BrdU+/GFP+/NeuN+** | 53.50% | 64.78% | 61.55% | 71.97% |
| **8 WPI** |  |  |  |  |
| **GFP+ Population: NeuN-** | 49.18% | 33.93% | 47.04% | 42.79% |
| **GFP+ Population: NeuN+** | 50.82% | 66.67% | 52.96% | 57.21% |
| **BrdU+/GFP+/NeuN-** | 22.15% | 21.00% | 31.43% | 20.15% |
| **BrdU+/GFP+/NeuN+** | 77.85% | 79.00% | 68.57% | 79.85% |

